# Homeostasis of columnar synchronization during cortical map formation

**DOI:** 10.1101/075341

**Authors:** Matthew T. Colonnese, Jing Shen

**Affiliations:** Department of Pharmacology and Physiology, Institute for Neuroscience, The George Washington University, 2300 I Street NW, Ross Hall 639, Washington DC 20037

## Abstract

Synchronous spontaneous activity is critical for circuit development. A key open question is to what degree is this synchronization models adult activity or is specifically tuned for circuit development. To address this we used multi-electrode array recordings of spontaneous activity in non-anesthetized neonatal mice to quantify firing rates, synchronization, binary spike-vectors and population-coupling of single-units throughout the period of map formation. Consistent with the first hypothesis, adult-like network interactions are established during the period of retinal waves, before the onset of vision and normal inhibition, and are largely conserved throughout juvenile ages. Significant differences from mature properties were limited to initial topographic map formation, when synchronization was lower than expected by chance, suggesting active decoupling in early networks. These findings suggest that developmental activity models adult synchronization, and that there is remarkable homeostasis of network properties throughout development, despite massive changes in the drive and circuit basis of cortical activity.

## INTRODUCTION

Connectivity during development is achieved by synapse formation under the control of molecular guidance cues, and modification of these synapses by neural activity (Katz and Shatz, 1996). In the immature cortex, the period of intense synaptogenesis and the initial formation of circuits and organized maps is also a time of dramatic shifts in the patterns of spontaneous and sensory evoked activity (Khazipov et al., 2013; Ackman and Crair, 2014). Early activity differs from adult in the amount of network silence, in the generators and frequency of network oscillations, and in the dynamics of the membrane potential during activation (Luhmann et al., 2016). However, to what degree these differences translate into unique network interactions during circuit formation is poorly understood, particularly among the layered columnar circuits of isocortex. Activity influences circuit formation by co-ordinating firing between pre and post-synaptic neurons resulting in synaptic stabilization or elimination (Zhang and Poo, 2001; Kirkby et al., 2013). Thus, the degree of synchronization is a critical characteristic that determines the mechanisms of activity dependent development (Butts and Kanold, 2010).

Multiple studies *in vivo* and *in vitro* have suggested that spontaneous activity during circuit formation is highly synchronized. Acute slices of cortex display waves of synchronized activity during a limited developmental period (Ben-Ari et al., 1989; Moody and Bosma, 2005; Allene et al., 2008). These unique activities are dependent on circuit configurations unique to different developmental periods (Dupont et al., 2006; Bonifazi et al., 2009). *In vivo*, early cortical activity is characterized by large bursts of rapid oscillations that synchronize multi-unit firing into brief windows of opportunity (Yang et al., 2009; Colonnese and Khazipov, 2010; Minlebaev et al., 2011; Brockmann et al., 2011). Calcium imaging of visual and somatosensory cortex *in vivo* shows that these early activities drive layer 2/3 firing that is hyper-synchronous relative to mature patterns (Golshani et al., 2009; Rochefort et al., 2009; Siegel et al., 2012), largely because early circuits lack an asynchronous ‘active’ state (Colonnese, 2014). A similar developmental trajectory occurs in slices of somatosensory and auditory cortex (Frye and MacLean, 2016).

These findings of apparent early hyper-synchrony support a ‘refinement’ model of circuit development. In this model, early hyper-connectivity caused by random synaptogenesis is replaced during a period of refinement with mature connections. In addition to the increased synchronization resulting from exuberant connectivity, circuit properties such as weak inhibition, excitatory GABA_A_ currents, long channel decay times, abundant electrical connectivity, and high neuron excitability should increase synchronization of local network firing (Blankenship and Feller, 2010; Cossart, 2011; Dehorter et al., 2012). By this model, immature hyper-synchrony drives the formation of low-frequency connectivity such as topography, before maturation of circuit properties renders the local network less synchronous, allowing network fractionation into local microcircuits, such as for orientation or direction selectivity in visual cortex (White and Fitzpatrick, 2007; Butts and Kanold, 2010). An opposed model is that early activity largely models adult activity, allowing for selectivity to emerge gradually and in parallel (Erwin and Miller, 1998; Crowley and Katz, 2002). This model predicts growing or static network synchronization. A corollary of this hypothesis is that neuronal and synaptic maturation does not desynchronize networks, but rather conserves network properties in the face of increasing synaptic density and excitability.

Clearly distinguishing these models is not currently possible as previous assays lack cellular resolution, measure activity only in single layers, or lack the temporal resolution to determine spike correlations within a cortical column. We therefore used multi-electrode array recordings combined with spike-sorting of units measured throughout the depth of individual cortical columns to measure synchronization in the developing visual cortex, a region for which the primary drivers of developmental activity and their role in circuit formation are largely known (Huberman et al., 2008; Ackman and Crair, 2014). During initial circuit formation (before eye-opening), activity in visual cortex is driven by spontaneous waves of activity in the retina (Ackman et al., 2012; Siegel et al., 2012; Kummer et al., 2016). These are amplified and shaped into oscillations by the unique properties of thalamic and cortical circuits (Weliky and Katz, 1999; Hanganu et al., 2006; Colonnese and Khazipov, 2010; Colonnese et al., 2010). These early network properties are replaced by the mature cortical circuit dynamics when true vision develops around eye-opening, as spontaneous activity increases and amplification of retinal input is eliminated by the development of feedforward inhibition (Rochefort et al., 2011; Colonnese, 2014; Hoy and Niell, 2015; Smith et al., 2015). We therefore performed multiple measures of synchronization in the network during the first three post-natal weeks, when cortical activity patterns are changing most rapidly. We find that adult-like network properties are established remarkably early and remain stable in the face of the extensive synaptic and circuit changes occurring over this period, supporting the conservative model of cortical columnar network development.

## RESULTS

We examined spontaneous activity in monocular visual cortex of unanesthetized, head-fixed mice. Recordings targeted 5 key developmental age groups: P6-7 & P8-9 during the period of cholinergic retinal waves, when topography and eye specificity is established; P10-11 during the period of glutamatergic retinal waves; P15-17 after eye-opening, during the pre-critical period (Smith and Trachtenberg, 2007) when cortical state modulation of spontaneous activity has emerged; and P24 during the critical period for ocular dominance plasticity when mature cortical dynamics are largely in place (Hoy and Niell, 2015). Before P6, spike-rates were not sufficient for clustering.

Network analysis was made by isolation of presumptive single-units from a single shank, dual column multi-electrode array placed perpendicular to the cortical layers, allowing for simultaneous recording from layer one though the top of layer six. Sorting using the masked EM algorithm(Rossant et al., 2016) was of similar quality between age groups (Fig. 1). Spike amplitudes in the youngest age group were lower, and clustered near threshold, but overall the similarity of spike wave-forms placed in the same cluster was comparable between age groups (Fig. 1 A&B). Refractory violations (inter-spike intervals < 2ms) were rarer in young animals, likely as a result of lower spike-rates (Fig. 2), making this a less reliable index of cluster quality in neonates. The number of well isolated (‘good’) clusters extracted from each animal increased rapidly with age (Fig. 1E). This was not a result of more clusters or spikes rejected in young animals. In fact, the proportion of total recorded spikes that were placed in good clusters was highest during the first two weeks (Fig. 1F). In general, while the lower size and reduced ability to use refractory violations to separate clusters increases the chances of multi-unit clusters, the likely reduced pickup volume (as evidenced by the total reduction in number of spikes and clusters) counteracts this effect. We are confident that clustering effectively enriches for single neurons in all age groups. Because analysis of fewer neurons could bias results between age groups, when appropriate, analyses were performed on a random selection of 12 units (the minimum number of good clusters isolated), calculating the relevant metrics, and repeating this process to generate a mean for the animal.

**Figure 1.**
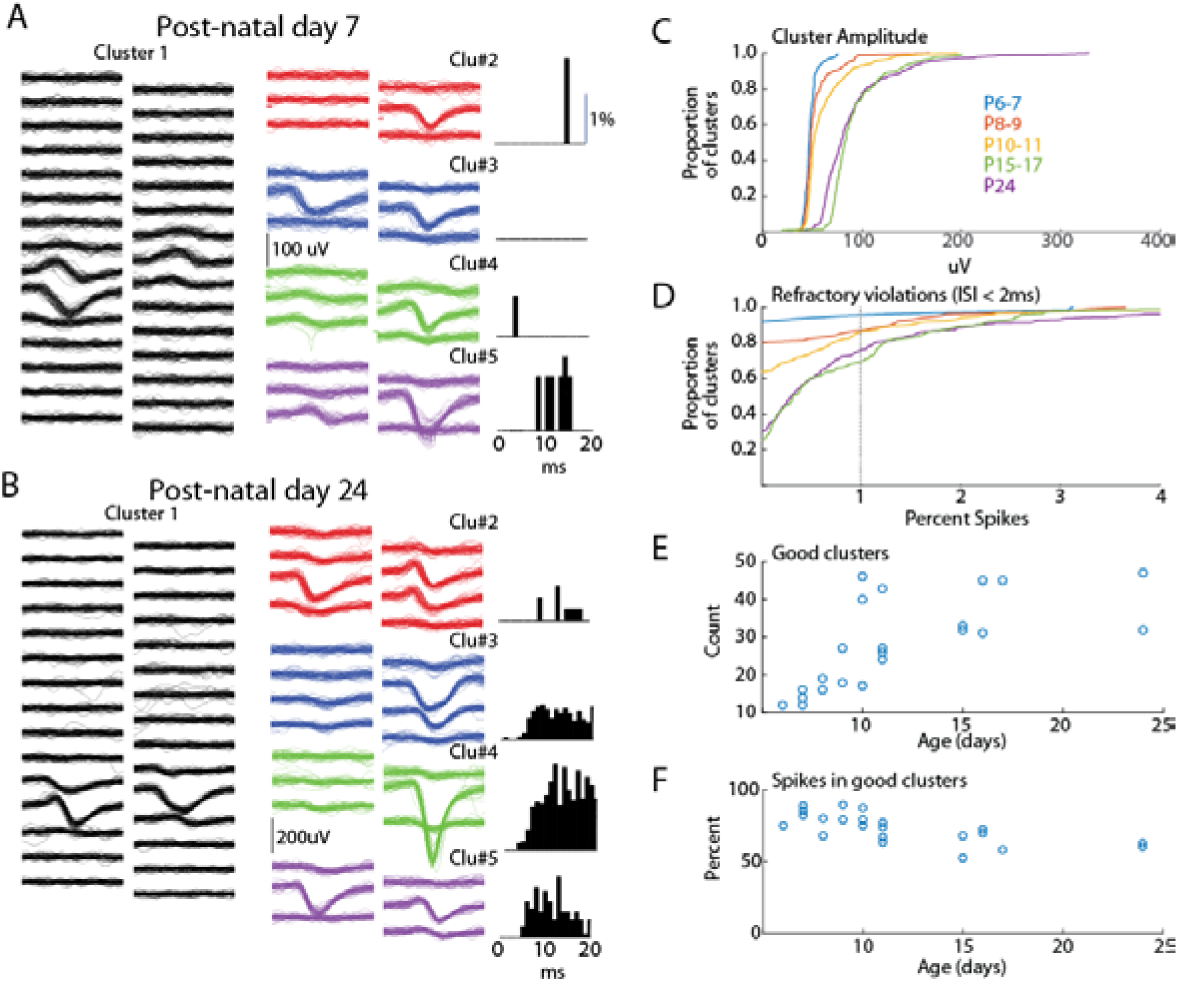
Spike-clustering in neonatal mice. **A.** Five representative clusters from same seven day old animal (P7). Each trace is 1ms, scale bar (100uV) applies to all traces. On left (Cluster 1) show 50 traces from 30 channels in the poly-2 array (50µm separation). 6 surrounding channels are shown for 4 additional representative clusters in the middle, and the associated inter-spike interval histogram for the same cluster is displayed at right. **B.** Representative clusters for P24 animal. **C.** Cumulative distribution of peak amplitude for all clusters (good and bad) in each age group used in the study. **D.** Cumulative distribution of the percent of refractory violations (interspike interval (ISI) < 2 ms) for all clusters. 1% violations was the threshold to reject clusters. Note fewer violations in young animals. **E.** Number of ‘good’ clusters isolated for each animal by age. **F.** Percentage of total recorded spikes that were placed in good clusters.

We first asked if age is associated with differences in the firing rates of individual neurons (Fig. 2). As previously shown for unanesthtized mice(Adelsberger et al., 2005; Ackman et al., 2012) and rats (Hanganu et al., 2006; Colonnese and Khazipov, 2010), activity during the first two weeks post-natal is dominated by lengthy periods of network silence (‘down-states’). The down-states are interrupted by periods of activation driven by spontaneous retinal waves (P2-9) and then by retinal waves as well activity generated spontaneously within cortex (Colonnese and Khazipov, 2010). Down-states lasting more than 200ms disappear around eye-opening in rats (Colonnese, 2014). Our unanethetized mice demonstrated a similar pattern, with single-unit activity almost completely restricted to 2-10s active periods occurring every 30-60 s. Extended periods of network silence were rare in the P15-17 or P24 groups (Fig. 2D).

**Figure 2.**
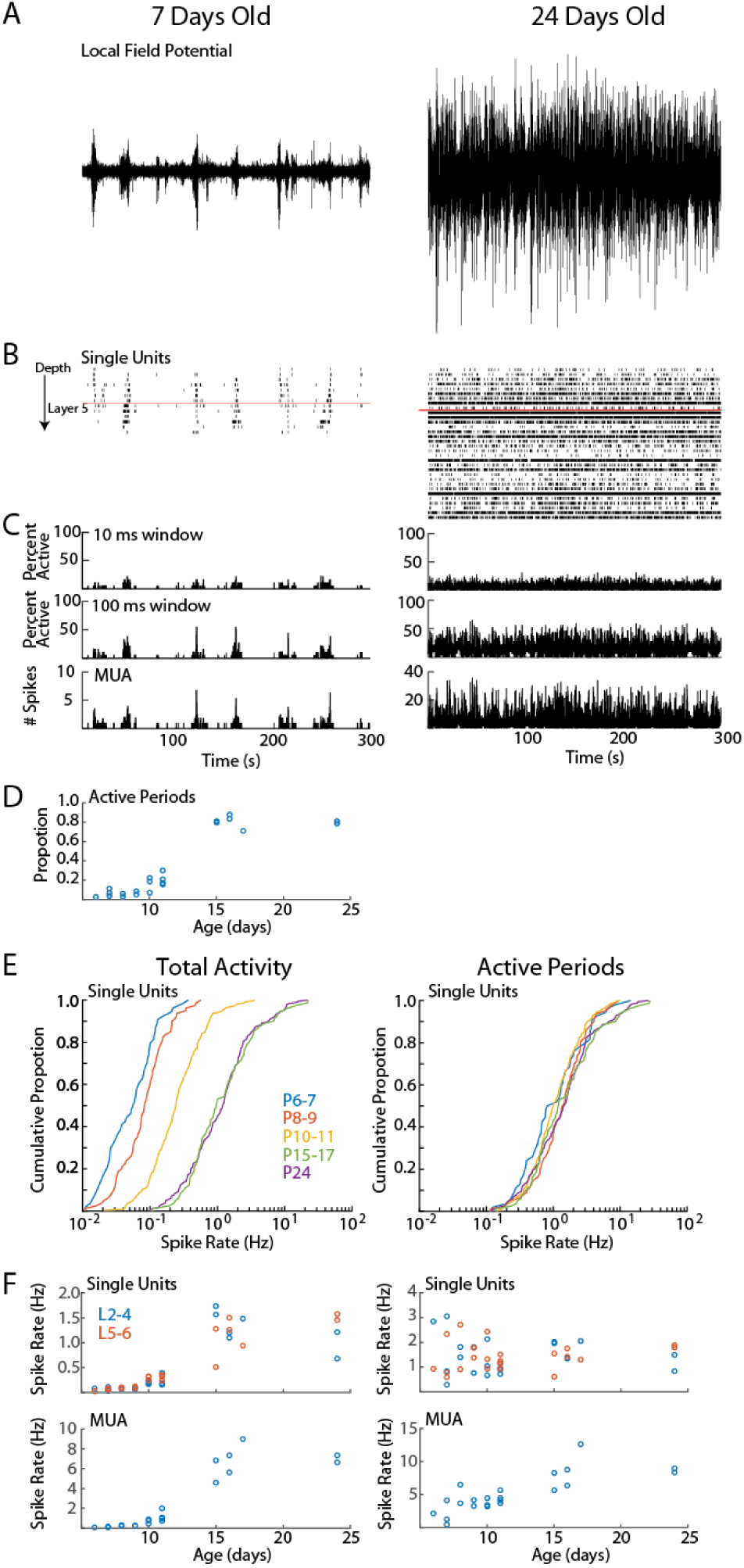
Cortical activity in neonatal rats is dominated by down-states. **A-C.** Representative spontaneous activity (300s) for a P7 and P24 rat. A. Local-field potential from layer 2/3. **B.** Raster plot of all good single-units arranged by depth. Red line shows top border of layer 5. **C.** Spike histograms showing percentage of single-units active in a 10ms (top) or 100ms (middle) window. At bottom the multi-unit activity (including spikes not sorted into good single-units) in 1 ms bins is displayed. **D.** Proportion of recording occupied by active periods (see Methods) for each animal, by age (ANOVA for effect of age (groups P6-7, 8-9, 10-11, 15-17, 24), df=4, F=188, p=10^-13^). **E**. Cumulative distribution histogram of spike-rate of all single-units in each age group. Total activity is shown on the left, firing rate during active periods (down-states removed) is on right. Developmental changes in single-unit firing rate are due to increased prevalence of down-states at these ages. **F.** Firing rates for single-units (above) and total multi-unit activity by age. Total activity (left) effect of age (L2-4 F=61.24, p=10^-9^; L5-6 F=38.46, p=10^-8^); MUA effect of age (F=60.72, p=10^-9^). Active periods only (right)(L2-4 F=0.92 p=0.47; L5-6 F=0.7, p=0.60); Active period only MUA (F=11.23, p=0.0001).

Total single-unit spike-rates at P6-7 and P8-9 were at least an order of magnitude lower than juvenile rates (Fig. 2E), with P10-11 rates intermediate. However if the down-states are removed and only periods of activation considered, then single-unit spike-rates were similar between ages, suggesting that active periods in young and old mice are similar, and that age differences in firing result from the prevalence of down-states during the first two post-natal weeks.

### Development of network synchronization and composition

Functional hyper-connectivity of inputs and/or local connections occurring either through increased connectivity or lack of desynchronizing inhibition is expected to result in activity events with high-participation rates, as has been observed by calcium imaging of layer 2/3 *in vivo* (Rochefort et al., 2009).

Because firing during early ages is largely restricted to the troughs of spindle-burst oscillations we first measured event participation in 20ms windows, approximately the window of firing during these early oscillations (Hanganu et al., 2006; Colonnese and Khazipov, 2010). To measure columnar event participation we calculated the percentage of single-units active (at least 1 spike) in each 20ms window. Event participation rates show clearly that early activity is not hyper-synchronous (Fig. 3A). In fact participation rates in the youngest group (P6-7) is similar to juvenile (P15-17 & P24), and activity is less synchronous at P8-9 and P10-11 before achieving stable values by P15 (Fig. 3C). Event participation is affected by spike-rate as well as spike timing. To control for the former we calculated the change in event probability after jittering spike times by a random amount ±1s, and calculating a deviation index (Prob – Prob_jitt_)/(Prob+Prob_jitt_). This analysis showed that event participation during the periods of retinal waves (P6-11) were actually lower than expected by chance, while the juvenile synchrony is slightly higher than expected (Fig. 3B,D).

**Figure 3.**
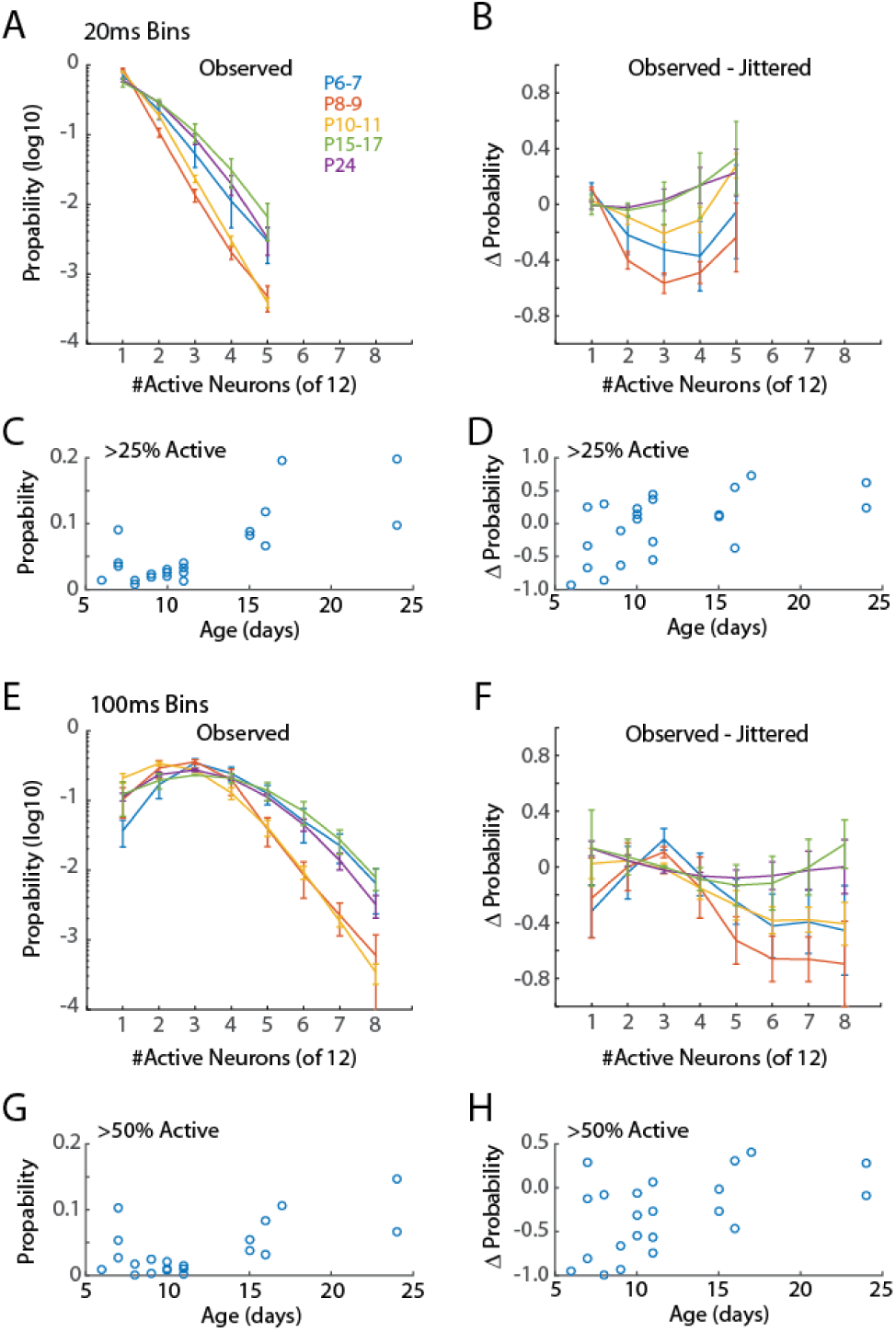
Synchronization of firing increases with age. A. Distribution of neuronal event size for 20ms windows. Points show mean and SEM of distributions for animals in each group. For each animal the probability of observing events of the indicated size in a 20ms window from a random assignment of 12 neurons is shown. Only active periods are considered. ANOVA for effect of synchronization (F=315.89 p=10^−49^), Age group (F=25.55, p=10^-13^) and interaction (F=4.41, p=10^-6^). **B.** Change in event size probability relative to jittered spike trains (Prob-Prob_jitt_)/(Prob+Prob_jitt_). Young animals (P6-11) have lower probabilities of synchronized events than expected by chance (ANOVA for synchronization F=5.14 p=0.0009; Age F=8.07, p=10^-5^; interaction F=1.16, p=0.32). **C.** Proportion of 20ms bins with more than 25% of single-units active for each animal by age. ANOVA for age group (F=7.33, p=0.0013). **D.** Change in probability (vs jittered) for >25% synchronization by age (F=2.38, p=0.09). **E.** As for *A* but 100ms window. Synchronization at P8-9 and P10-11, but not P6-7, is lower than juvenile ages. ANOVA for effect of synchronization (F=82.69 p=10^-44^), Age (F=15.55, p=10^-10^), and interaction (F=5.21, p=10^-11^). Highly synchronized events P6-11 are less likely than chance. **F.** As *B* but 100ms window (Synchronization F=4.53 p=0.00014; Age F=6.93, p=10^-5^; interaction F=0.92, p=0.59). **G.** As *C* but probability of events with >= 50% synchronization (F=4.04, p=0.0175) of units are shown. **H.** As for *D* but events >= 50% synchronization (F=2.13, p=0.122).

Currents in young neurons have longer decay times, potentially increasing the integration time and tolerance for synchrony. We therefore examined a longer time window for neural events (100ms) which encompasses a complete cycle of the early oscillations. Developmental patterns of event participation at 100ms were largely similar to 20ms windows (Fig. 3E-H). Thus, regardless of window size, event participation rates of neonates are lower than those of juveniles and lower even than expected by chance given neuronal firing-rates. This suggests that activity in neonatal cortex is actively decoupled.

Our event participation data reject the hypothesis that mature neuronal ensembles are formed by fractioning larger ensembles, but obscure the specifics of which neurons fire together. To understand the development of neural ensembles we used an analysis of the occurrence of unique binary spike vectors, or ‘neural words’ (Fiser et al., 2004) (Fig. 4A). The occurrence of specific words is tested against the distribution expected by random firing, modeled here by random jittering the occurrence of spikes ±1s. We examined word distributions for words 3 neurons or longer in 20ms windows.

**Figure 4.**
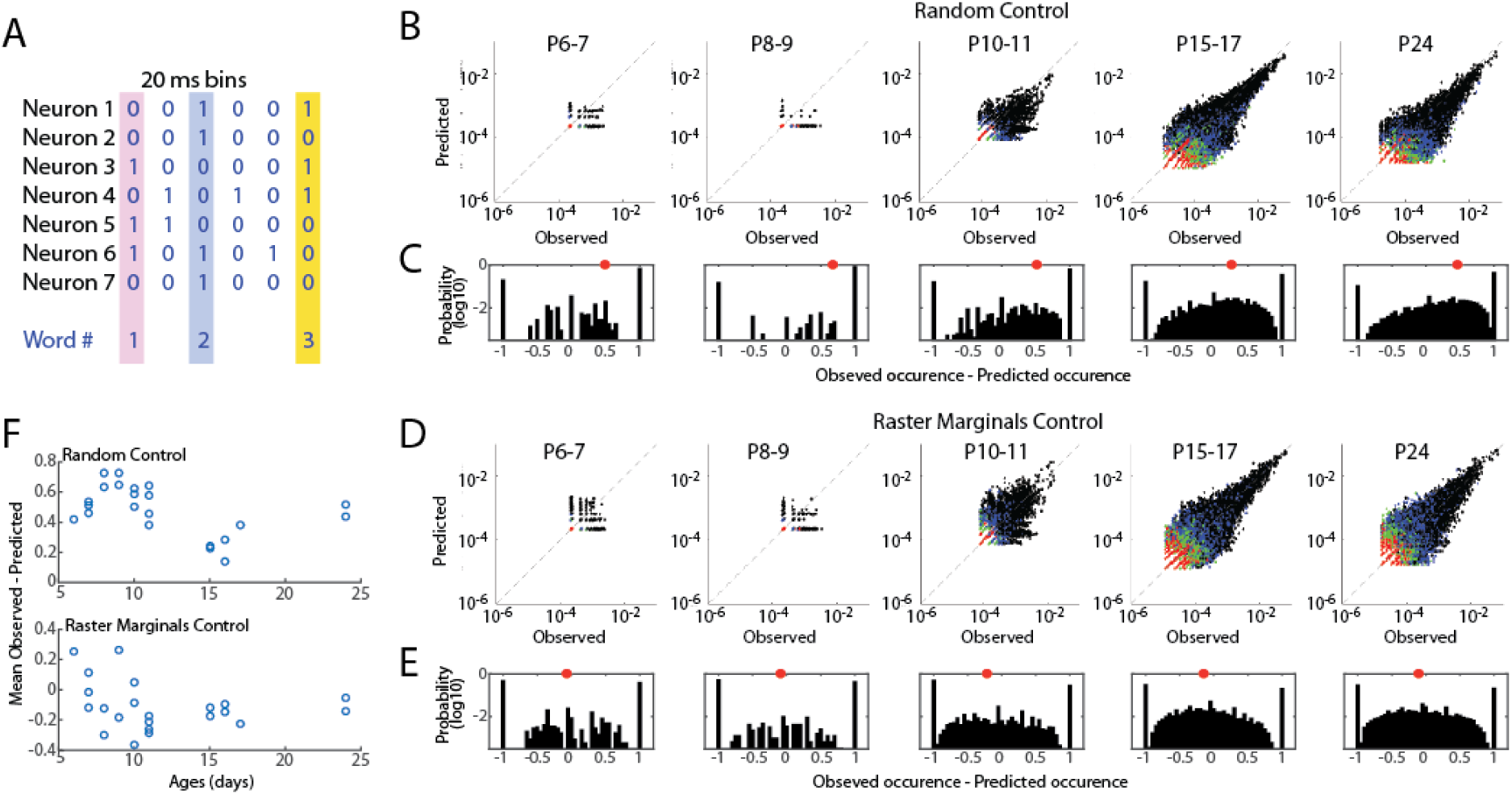
Developmental conservation of neuronal assembly properties. **A.** Measurement of neuronal assemblies as unique ‘words’ consisting of binary firing vectors for 20ms windows. Only assemblies of 3+ units are considered. **B.** Probabilities of observed word occurrence vs. occurrence predicted by random jittering of spikes. Graphs show all words in each age group. Black dots are 3 neuron words, blue 4, green 5, red >=6. **C.** Probability distribution for observed – predicted occurrence of all words in each age group. 1 means that word occurred only in observed data; -1 means it occurred only in jittered data. Distribution is right shifted at all ages, indicating greater occurrence of words than expected by chance throughout development. Red dot shows mean of distribution. **D.** As *B* but predicted probability is calculated by Raster Marginal method (Okun et al., 2012) which controls for changes in network properties controlling the timing of spikes. **E.** As *C* but for Raster Marginals control. **F.** Mean difference of word distributions from random (top) or raster marginal (bottom) model for each animal by age. ANOVA for effect of age group (Random F=17.45, p=10^-5^; Marginal F=1.14, p=0.374).

As expected given the larger number of good clusters in older animals, the total number of observed words increased with development, however word occurrence was greater than expected for random activity at all ages (Fig. 4B-C). The mean of the observed probability-predicted probability (by chance) was larger in in P6-11 animals than P15-17 & P24 (Fig. 4F). This relationship could indicate that connectivity in the young network is more strongly non-random than juvenile, or that network properties which synchronize spike-timing regardless of ensemble participation are stronger in young animals (Okun et al., 2012), which is consistent with the presence of spindle-burst oscillations during this time period. To distinguish these possibilities we predicted word occurrence distributions using the Raster Marginal method (Okun et al., 2012), which swaps spikes between clusters, thereby keeping intact temporal restraints on spike-timing but randomizing occurrences within clusters. Mean observed-predicted occurrence using the marginal control was near zero for all ages, and not different between age groups (Fig. 4D-F), suggesting that at all ages word distributions are similar to that expected of a random network in which spike timing is constrained by synchronizing mechanisms.

### Neural firing rate correlations increase during development

Spike-rate correlations reflect neural connectivity filtered through cellular, synaptic and circuit properties influencing synchronization, particularly inhibition (Renart et al., 2010; Helias et al., 2014). Ineffective inhibition, neuronal hyper-excitability and hyper-connectivity would be expected to increase spike-rate correlations in infant cortex, as has been observed for calcium signal in somatosensory cortex (Golshani et al., 2009). To test this we calculated the distributions of pair-wise spike-rate correlations for all good units in an animal. Spike-rate correlations were evaluated by quantifying spike-rate comodulation within a rapid time window (*T*=20ms full width at half amplitude Gaussian) corrected for slow modulation of firing rates (*J*=80ms)((Renart et al., 2010). Pairs were also subdivided into superficial (L2-4) and deep (L5-6) to assay the development of local vs total synchronization. Because the presence of down-states, which increase correlations by enforcing silence in almost all neurons, changes dramatically during development, we examined correlation distributions for total activity as well as when restricted to active periods.

The pattern of spike correlation distribution for total activity was grossly similar at all ages (Fig. 5). The large majority of pairs had correlations near zero, but the distribution evidenced a right-ward shift toward a larger proportion of correlated neuron pairs than expected by chance (jittered spikes) that grew stronger with age. As a result the mean spike correlation of each animal increased when measured between all neurons as well when restricted to deep neurons, but this change was not significant for superficial neurons (Fig. 5B). Limiting analysis to active periods caused a left-ward shift in the correlation distribution at all ages. This shift was much larger for neonatal animals, even resulting in mean correlations below zero for the P6-7 and P8-9 groups. Thus at these youngest ages, firing rates are less correlated on a short time-scale (20ms) than expected by chance, suggesting inhibitory or other desynchronizing element are powerful even during early development. As a result of the negative mean correlations in neonates, developmental increases in mean correlation were significant for total, superficial and deep neurons (Fig. 5B).

Calcium imaging studies have described developmental desynchronization of layer 2/3 activity (Golshani et al., 2009; Rochefort et al., 2009), opposite from the increasing synchronization we observed here. While multiple factors could contribute to this difference, the effective integration window for firing is a prominent difference between our spike-rate correlations and calcium imaging. To examine the role of integration window on mean spike-rate correlations, we systematically varied *T* (with *J* also increasing at 4\**T*). This showed a strong, age dependence of correlation on integration window (Fig. 5C). For total activity, as integration windows approach 400ms the developmental relationships between neonatal and juvenile animals reverse, with activity P6-11 becoming more synchronous than P15-17 & P24. When correlations were limited to active periods, integration window does not have as dramatic an effect. However, negative mean correlations present in neonates were only present when *T* was less than 300 ms. To determine if this dependence of integration window was the result of changing *T* or *J*, we varied *J* while keeping *T* constant. Correlations were largely unchanged out to a J of 3s (data not shown), beyond the duration of a single retinal wave (Blankenship and Feller, 2010; Ackman et al., 2012). Finally, because maximal negative mean correlation is inversely proportional to number of neurons, we recalculated pair-wise correlations for N=12 neurons in all groups (data not shown). Mean correlations were not significantly different in this N limited case, showing that the growth of correlation with age is a true effect of development, not of the number of neurons isolated by spike-sorting.

**Figure 5.**
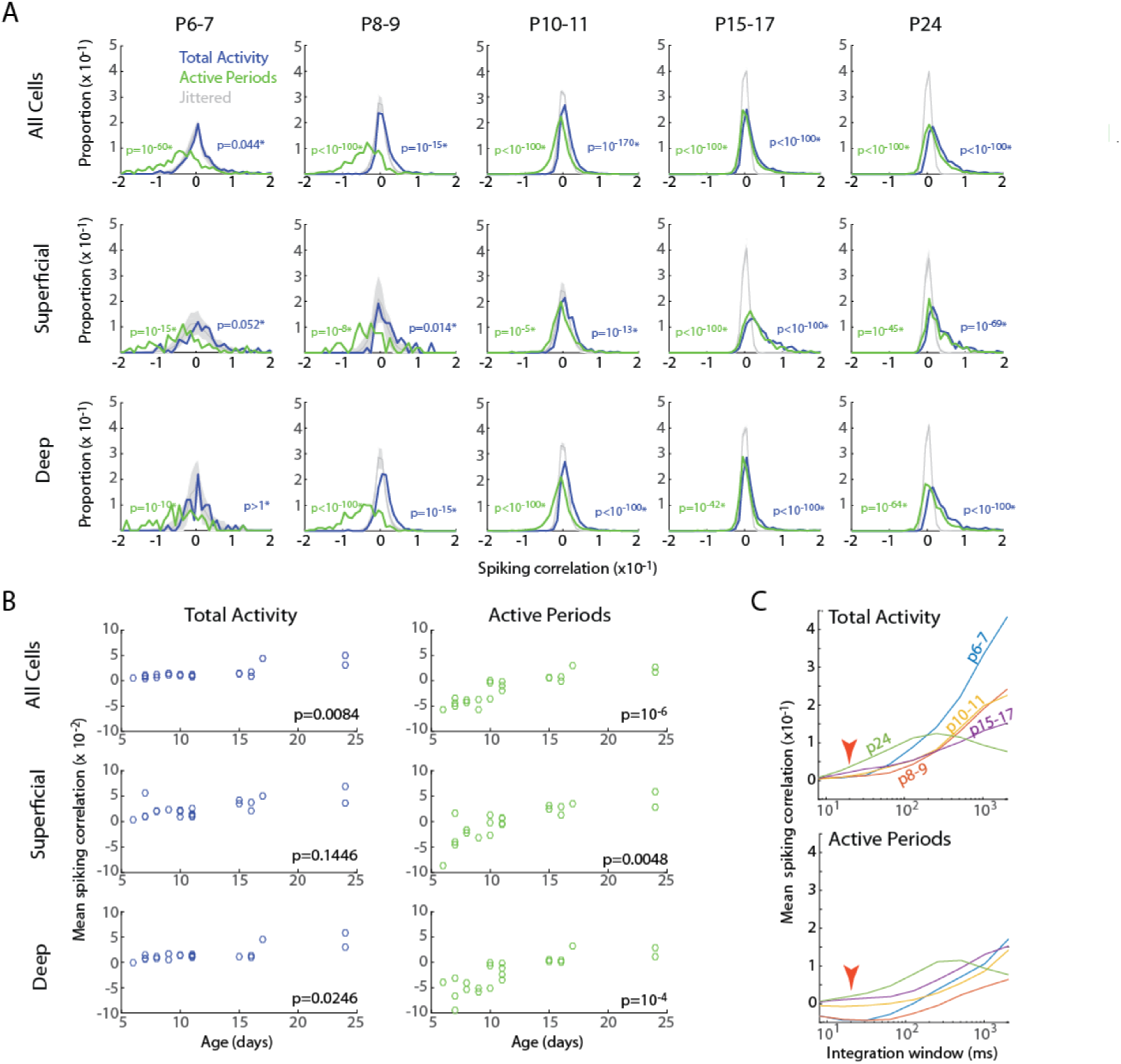
Developmental increase in pairwise spiking correlations. **A.** Distribution of pair-wise firing rate correlations for all neuron pairs in each age group. Blue line shows correlations for total activity including down-states; green line shows correlations when analysis is limited to active periods. Grey shading shows 95% confidence interval of jittered correlations. P-values are results of K-S test for difference from jittered distribution (*indicates p-values were multiplied by 15 to account for multiple comparisons). Top row shows all pairs. Middle row is distribution for superficial vs superficial neurons only (L2-4). Bottom row for deep vs deep neurons only (L5-6). **B.** Mean correlations for each animal by age for the Total (left) and limited to Active periods (right) for each of the neuron populations. Listed p-values are for ANOVA effect of age group. **C.** Effect of integration window (SD of Gaussian filter) on spiking correlations. Red arrow shows integration window used for figure (20ms). Developmental increase in mean correlations for total activity reverses at large integration windows.

In sum, while the pair-wise firing rate correlations are sensitive to integration window, within physiologically relevant time intervals (10-300ms), they are robust and consistent with increasing synchronization of the neural activity during development. Combined with event participation, firing rate correlations suggest that early networks contain decorrelating influences that keep synchronization below that expected by chance given firing rates and patterns.

### Characteristics of local network integration are established early in development

Neurons vary in their local vs distal connectivity, a feature that is correlated with the degree to which their firing is coupled to mean firing rates in the local network, called ‘population coupling’ (Okun et al., 2015). We hypothesized that hyper-connectivity during early development would result in increased population coupling and fewer neurons with activity that is independent of local firing (so called ‘soloists’). To test this we calculated the spike-triggered multi-unit activity for each good unit. Triggered spike-rates are converted to standardized ‘population coupling’ by normalization to the same measure for spikes jittered using the Raster Marginal method, thereby constructing a mean coupling for the animal that can be used to normalize between groups.

**Figure 6.**
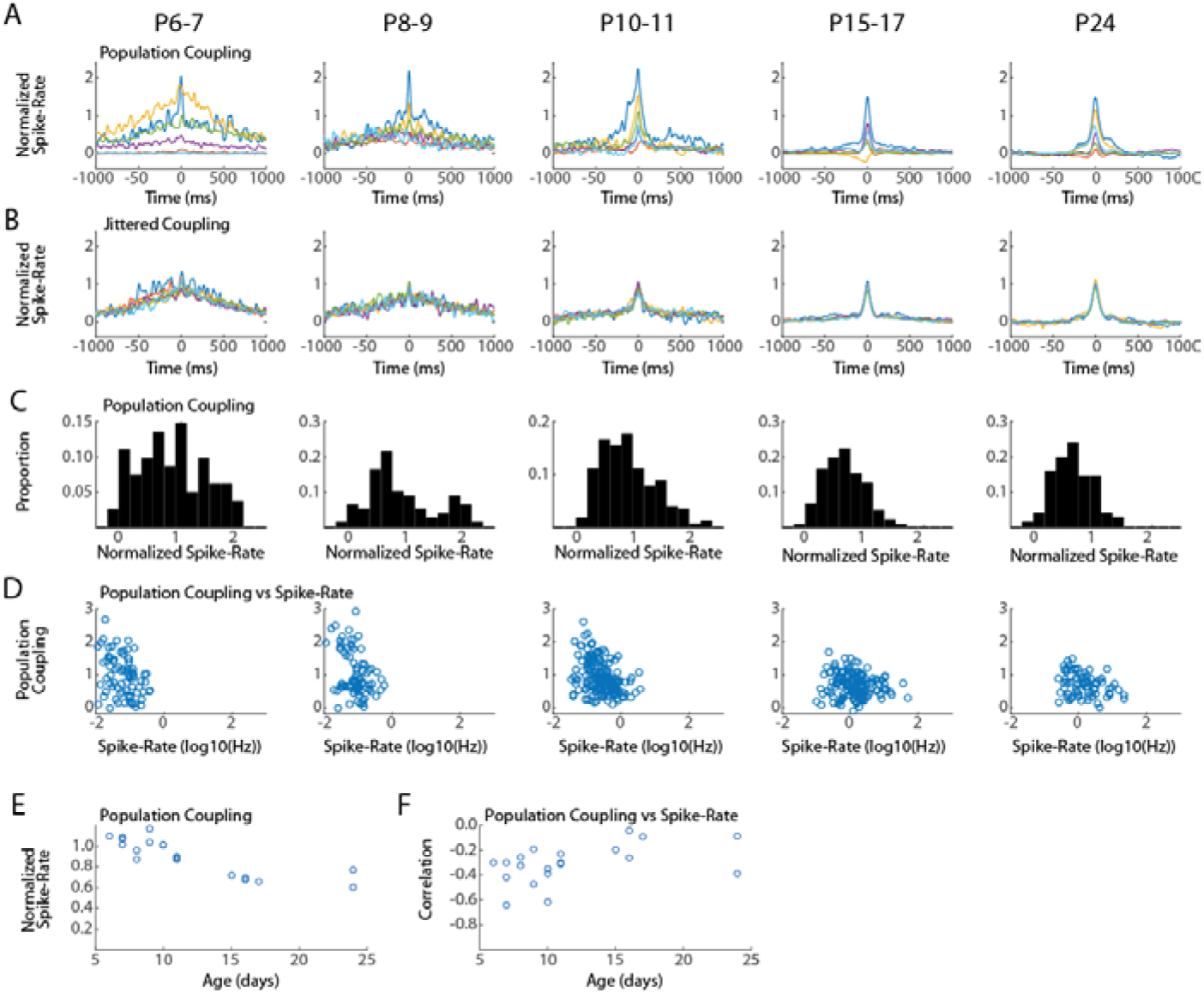
Conservation of local network integration during development. Population coupling (normalized spike-triggered multi-unit firing rate) is correlated with local connectivity in adults (Okun et al., 2015). **A.** Representative population couplings for five neurons from a single animal in each age group. Each animal has neurons with diverse coupling, though the temporal specificity of the coupling increases with age. **B.** Coupling for the same neurons after jittering. **C.** Distribution of population coupling for all single-units in the age group. **D.** Spike-rate plotted against population coupling for all neurons in each age group. Spike-rate changes between ages do not strongly affect results, though presence of very low spike-rates appears required for high population coupling. **E.** Mean population coupling for each animal by age. ANOVA for effect of age group (F=10.24, p=0.0003). **F.** Mean correlation of coupling vs spike-rate for each animal by age. ANOVA for effect of age group (F=1.82, p=0.1748).

The temporal characteristics and signal-to-noise of population coupling changed dramatically between P6-7 and P15-17, but all ages showed dramatic variance in the absolute degree of coupling. Every animal had neurons with strong coupling as well as neurons with little or even negative coupling (Fig. 6A). The total distribution of normalized spike-rates showed a similar pattern of diverse coupling in each age group, with a peak between 0.5 and 1 (Fig. 6C). The two youngest age groups (P6-7 & P8-9) had an additional population of highly coupled neurons that was not apparent in the older ages. As a result, mean coupling was reduced with age (Fig. 6E). In adults, low population coupling is not correlated with low-spike-rates (Okun et al., 2015), but this relationship may not hold for young animals. We therefore examined the correlation between population coupling and spike-rate for individual neurons in each age group (Fig. 6D). At all ages low-population coupling occurred for all spike-rates, and the two measures were negatively correlated for all age groups (Fig. 6F), suggesting that same processes underlie the diversity of population coupling throughout development.

In total our results show that while population coupling is elevated for a sub-population of low-firing neurons during early development, neurons with weak and strong population coupling exist even in the youngest networks. These results suggest that that a neuron’s relative local integration is established early and maintained throughout development.

## DISCUSSION

Here we used multi-electrode array based spike-sorting in very young, head-fixed mice to show that adult-like network properties of synchronization, population coupling and even firing rates are established very early, prior to vision, when intracortical and thalamocortical circuits are forming and spontaneous retinal activity dominates cortical activation (Ackman and Crair, 2014). Once established, network properties are conserved despite critical changes in the patterns and origins of activity, circuit development—including functional inhibition--and the onset of vision and critical periods for plasticity (Huberman et al., 2008). Our results support the hypothesis that early spontaneous activity provides a model for adult activity, and that maturation of inhibition and other synaptic properties provides homeostatic control to maintain synchronization in the face of increasing excitation.

### Electrophysiology and imaging approaches in network analysis

Our results differ from *in vivo* calcium imaging studies, which consistently indicate a decrease in correlation between horizontally separated layer 2/3 neurons, both in in somatosensory (Golshani et al., 2009) and visual (Rochefort et al., 2009; Siegel et al., 2012) cortex. The primary contributor to this difference is likely the time course over which synchronization is measured. In our data, integrating spike-rates over large windows on the timescale of calcium indicators selectively increased the pairwise correlations of young neurons (Fig. 5C). Current-clamp recordings suggest that early hyper-synchronization observed in imaging is due to increased firing probability during periods of global activation (Golshani et al., 2009; Rochefort et al., 2009; Colonnese, 2014), and not co-participation of neurons in local ensembles, which appears to grow (Berkes et al., 2011) or remain similar (Fig. 3) during development.

One important caveat to the current results is that with the increased temporal and single-spike resolution of electrophysiology comes the inherent ambiguity of spike-sorting. The smaller transmembrane currents for action potentials in the young neurons make them less likely to be detected on multiple electrodes, makes waveforms closer to threshold, and reduces inter-neuron variability of waveforms, all of which potentially reduce the ability to sort. Current spike-sorting quality metrics emphasize separation and reduction of over-splitting (Hill et al., 2011) and the application of these metrics to the high-dimensional space of large arrays is not standardized (Rossant et al., 2016; Harris et al., 2016). Confirmation that the spikes in a single cluster originate from a single neuron relies on visual confirmation of waveform and elimination of clusters with high rates of refractory period violation. The low spike-rates in young animals should cause an increase in false negatives (failure to reject clusters with 2+ neurons). Thus while spike waveform consistency was similar between ages (Fig. 1), it remains possible that more “good” clusters in young animals contain multiple neurons. The developmental growth in correlations is unlikely to result from poor sorting however, as it is also evident in multi-unit activity in rat cortex measured at similar ages (Berzhanskaya et al., 2016).

While our mice were recorded un-anesthetized, which is critical because even sub-analgesic levels of anesthesia eliminate early activities (Adelsberger et al., 2005; Colonnese and Khazipov, 2010; Ackman et al., 2012; Sitdikova et al., 2014; Lebedeva et al., 2016), our data may not perfectly reflect natural activity patterns. In addition to the invasive nature of electrical recordings and tissue damage it causes, the animals are head-fixed potentially causing stress-related changes in synaptic transmission (Inoue et al., 2013) and abnormal release of neuromodulators which can influence cortical activity (Hanganu et al., 2007; Janiesch et al., 2011). In addition, the anesthesia used to place head-restraint can kill neurons (Istaphanous and Loepke, 2009).

### Constructionism vs refinement

Our results support a ‘constructivist’ or elaborative model of intra-columnar cortical circuit development in which correct connections, informed by guidance molecules and confirmed by activity, are largely made early, without large scale elimination of incorrect connections (Quartz and Sejnowski, 1997; Katz and Crowley, 2002). We observed no reduction in event participation, word distributions, pair-wise correlations, or population coupling which would be expected if the network experienced a period of widespread hyper-connectivity followed by synapse elimination. This does not mean that inappropriate synapses are not formed and eliminated. Rather, it suggests that inappropriate synapses are a minority of total synapses, and play little functional role in driving early activity. Map formation in the cortical column may be different from sub-cortical regions such as thalamus and superior colliculus that are the initial recipients of topographic connections. These regions do incur periods of exuberant synaptic connectivity and elimination (Chen and Regehr, 2000; Lu and Constantine-Paton, 2004; Ziburkus and Guido, 2006; Huberman et al., 2008), though even in these regions activity is highly localized and functional topography is apparent even from an early age (Akerman et al., 2002; Ackman et al., 2012). Our results are consistent with multiple findings that thalamocortical and cortico-cortical connectivity is refined very early (Katz and Crowley, 2002; Ko et al., 2013; Yang et al., 2013). Our results expand upon these by showing that even within topographically aligned columns, functional connectivity is sparce during initial map formation. Early and consistent refinement may be specific to intra-columnar circuits, which reflect a single topographic location, while horizontal connectivity may be subject to different rules. Though studies have observed both developmental increases and decreases in horizontal synchronization (Callaway and Katz, 1990; Fiser et al., 2004; Minlebaev et al., 2011).

One of our more unexpected findings is that variance in population coupling is present from as early as intra-cortical synapses are present, and can drive activity locally (Blue and Parnavelas, 1983; Valiullina et al., 2016). While we cannot prove that early ‘soloists’, neurons with low population coupling (Okun et al., 2015), are the same neurons that become adult soloists, our data suggest that this identity is set early, even before many of the long-range connections that will drive soloists are formed.

### Mechanism of activity dependent development

What do our results suggest about the mechanisms of activity dependent-circuit formation? The temporal fidelity of retinal waves is insufficient to explain topographic map formation using spike-timing plasticity alone (Bennett and Bair, 2015). Burst based synaptic plasticity provides another mechanism of refinement, and has been identified in thalamus and superior colliculus (Butts et al., 2007; Shah and Crair, 2008) but not cortex. Synaptic weakening occurs at synapses that do not consistently fire during the same retinal wave as the dominant synapses (Winnubst et al., 2015), further suggesting plasticity rules using long-duration time courses are important in cortex. We show, however, that the neonatal cortex can synchronize activity among refined ensembles at fast time courses. Retinal waves generate spindle-burst oscillations, which synchronize firing in ~20ms windows during their troughs (Hanganu et al., 2006; Colonnese and Khazipov, 2010). Such exquisite early synchronization likely uses transitory ‘booster’ circuits such as subplate (Tolner et al., 2012), or corticothalamic feedback excitation (Murata and Colonnese, 2016). In somatosensory cortex, these oscillations (as well as faster ‘early gamma oscillations’) are generated in thalamus (Yang et al., 2016), and frequencies above 10 Hz are critical to induce synaptic potentiation (Minlebaev et al., 2011). The present work extends these findings by showing that neurons do not fire during each phase of the spindle-burst oscillation, but rather each phase triggers separate local microcircuits sequentially, similar to beta-gamma oscillations in adult cortex (Harris et al., 2003). By showing that spike-correlations on a rapid timescale are adult-like in young cortex we further provide evidence that spike-time dependent plasticity on a rapid time scale is a potential mechanism of synaptic refinement during development.

While much is still poorly understood about the process leading to the formation of refined cortical ensembles, our data clearly indicate they do not emerge from larger, less refined functionally connected groups of neurons. In fact, in the youngest animals (P6-7 & P8-9) neuronal firing is actually less synchronous than would be expected by chance. This was true for both the participation rates as well as pair-wise correlations. A similar trend was observed in the neuronal word distributions, but it was not significant. Cortiocortical connectivity, both electrical and chemical, is very low at these ages (Yu et al., 2012), so low correlations would not be surprising. However, the negative correlations require a desynchronizing element to decorrelate activity driven by the massively synchronous spindle-burst oscillations coming from thalamus (Helias et al., 2014). During this limited early period, thalamic axons synapse on inhibitory subplate neurons (Kanold and Luhmann, 2010) as well as somatostatin neurons in layer 5 (Marques-Smith et al., 2016; Tuncdemir et al., 2016) before shifting to their adult targets, providing one possible mechanism of inhibitory desynchronization. An implication of this anti-correlation is that the net effect of correlation based plasticity should be toward elimination of new synapses, a phenotype observed in superior colliculus for the same ages (Colonnese and Constantine-Paton 2006).

### Circuits, synapses and synchronization: the more they change the more they stay the same

Developing cortical networks undergo remarkable changes in the amount and pattern of neural activity (Khazipov et al., 2013). In visual cortex, a large majority of the changes occur in rapid succession around eye-opening, though they are not strongly dependent on patterned vision. At this time immature spindle-burst synchronization of neural firing ends, cortical waking and sleep states emerge, thalamic amplification of retinal input is down-regulated, and the capacity of the circuit to follow relevant high frequencies emerges (Colonnese et al., 2010; Rochefort et al., 2011; Colonnese, 2014; Hoy and Niell, 2015). Somatosensory cortex makes a similar shift, though 4 days earlier, perhaps because whisking starts earlier than eye-opening (Colonnese et al., 2010; Minlebaev et al., 2011). Human infants undergo a similar shift before 2-4 weeks before term (Tolonen et al., 2007; Colonnese et al., 2010; Chipaux et al., 2013). The synaptic and network mechanisms of this shift are unknown, though they likely involve increased action potential threshold, development of ascending neuromodulators and functional integration of GABAergic interneuron subtypes, particularly those mediating fast-feedforward inhibition (Luhmann and Prince, 1991; Daw et al., 2007; Golshani et al., 2009; Colonnese, 2014). Inhibition is a controlling factor in many developmental transitions, particularly the onset of ocular dominance plasticity, leading to the suggestion they are ‘master’ regulators of development, transforming activity in order to switch function (Le Magueresse and Monyer, 2013). We observed remarkable stability of the network properties between P10-11 and P15-16, ages between which feedforward inhibition develops in visual cortex (Colonnese, 2014). Thus, our results suggest an alternate framework, which is that inhibitory (among other) development occurs to maintain firing-rate and synchrony homeostasis in the face of increasing synaptic density and its inherent excitability (Hengen et al., 2013). By this model, interneuron integration occurs not as a developmental program to transform activity, but as a bulwark against increasing activity resulting from excitatory synaptogenesis. Changes in the pattern of neuronal oscillations occurring at the same time may in fact be side-effects of the circuit changes obscuring the deeper similarity between early and late ages.

One conclusion of this homeostasis of network synchronization is that retinal-wave activity (which dominates P6-11 firing) does not drive unique early ensembles of high-synchronous firing designed to induce wiring of a single column, a conclusion also observed by L2/3 calcium imaging (Siegel et al., 2012), but rather to model adult cortical activity. It should be noted that the maintenance of correlational structure, does not imply that individual neurons maintain connections across development. In fact, mature local ensembles form by rearranging specific connections while maintaining the same total connectivity after eye-opening (Ko et al., 2013). Our result show that despite large scale changes in the factors that regulate synchronization in adults, network properties in young networks are maintained so that the firing correlations caused by early connectivity can be read out and modified to drive circuit ‘refinement’.

## MATERIALS AND METHODS

### Animal care

Animal care and procedures were in accordance with *The Guide for the Care and Use of Laboratory Animals* (NIH), and approved by the Institutional Animal Care and Use Committee at The George Washington University. Postnatal day (P)0 is the day of birth. C57BL/6 were obtained from Hilltop Lab Animals (Scottsdale, PA) as timed pregnant females, and kept in a designated, temperature and humidity-controlled room on 12/12 light/dark cycle.

### In vivo electrophysiology

Carprofen (20 mg/kg) saline was injected 1 hour prior to surgery to reduce pain and inflammation. Surgical anesthesia was induced with 3% isoflurane vaporized in 100% O_2_, verified by tail-pinch, then reduced to 1.5-3% as needed by monitoring breathing rate. A vented warming table (36°C, VetEquip, Livermore CA) provided thermoreplacement. For attachment of the head-fixation apparatus, the scalp was excised to expose the skull, neck muscles were detached from occipital bone, and the membranes were removed from the surface of the skull. Topical analgesic was applied to the incision of animals older than P8 (2.5% lidocaine/prilocaine mix, Hi-Tech Pharmacy Co., Inc., Amityville NY). Application to younger animals was lethal and discontinued. The head-fixation apparatus was attached to the skull with grip cement (Dentsply, Milford DE) over Vetbond™ tissue adhesive (3M). Fixation bar consisted of a custom manufactured rectangular aluminum plate with a central hole for access to the skull. After placement, the animal was maintained with 0.5-1% isoflurane until the dental cement cured, after which point it was allowed to recover from anesthesia on the warmed table.

For recording, animals were head-fixed and body movements were restricted by placement in a padded tube. Body temperature was monitored via thermometer placed under the abdomen, and maintained above 33C and below 36C via thermocoupled heating pad (FHC, Bowden ME). Body motion was monitored with a piezoelectric device placed below the restraint tube. For electrode access, a craniotomy was performed thinning the skull if necessary and resecting small bone flaps, to produce a small opening (~150-300 µm diameter). Primary visual cortex was targeted by regression of adult brain lambda-bregma distances: 1.5-2.5mm lateral and 0.0-0.5 mm rostral to lambda. All recordings were made using a single shank, 32 channel array arranged in two parallel lines of contacts (A1x32-Poly2-5mm-50s-177, NeuroNexus Technologies, Ann Arbor MI). The electrode penetrated the brain orthogonally to the surface and advanced to a depth of 750-1000 µm using micro-manipulator (Narishige, Japan) until the top channels touched the dura. Isoflurane was withdrawn and the animal was allowed to acclimate inside the setup for at least 50 min prior to recording. Recording sessions lasted 80-120 min after acclimation. The spontaneous activity reported here was recorded after 30 min of quiet rest in the dark and 20 min of visual stimulation. Recording lasted 30 min. All recording was performed in the dark (<0.01 Lumens). In animals older than P8, recording localization in monocular V1 was confirmed by the presence of a contralateral visual response to whole-field light flash that had earliest response in layer 4. Ipsilateral visual LFP responses less than 10% of the contralateral response were also required. All animals were sacrificed by anesthetic overdose followed by decapitation. Brains were immersion fixed in 4% paraformaldehyde for confirmation of electrode location.

### Data acquisition and analysis

Data was acquired at 32 kHz using the Digital Lynx SX acquisition system and Cheetah version 5.6.0 (Neuralynx, Inc., Bozeman, MT). Signals were band-passed 0.1 Hz – 9 kHz and referenced to a sub-cortical contact in at the bottom of the array. Analysis utilized custom MATLAB (MathWorks, Natick, MA) routines and the open-source *Klusta*suite for spike isolation, clustering and manual curation (Rossant et al., 2016). A spike isolation strong threshold of 6SD and weak threshold of 3SD were needed to minimize low amplitude unclusterable spikes. Initial clustering was evaluated for merging or splitting of clusters based on visual waveform analysis and similarity. After eliminating clusters composed of noise, clusters with no modulation of autocorrelation near 0ms and/or clear superposition of 2 or more distinct waveforms that could not be separated were marked as multi-unit clusters. The remaining, potentially single-unit, clusters were evaluated for inclusion as good single-units if the interspike interval refractory violations (<2ms) accounted for less than 1% of spikes. All clusters were assigned a primary contact localization by determining the minima of the mean waveform. Spike-time is was assigned by rounding the time peak time to the nearest ms. A total multi-unit (tMUA) raster was created by summing spike occurrences of all multi and single-unit clusters. The layer identity of each channel was made relative to L4 which was identified in an age specific manner. After the emergence of visual responses on P8, L4 was identified as the channel with the shortest latency 300-500µm below the surface. For P4-P7, which lack visual response, L4 was identified from spontaneous spindle-bursts as the lowest channel with visible rapid oscillations in the LFP (Colonnese and Khazipov 2010).

Inactive periods (down-states) were identified by the method of (Renart et al., 2010). The tMUA signal was convolved with a Gaussian kernel of SD 20ms and periods where this convolved signal was less than 10% of the animal’s peak tMUA and where tMUA interspike interval was less than 50ms were considered active periods. Active periods shorter than 50ms were eliminated. Event size was calculated by first selecting a random subsample of 12 single-units and a matrix consisting of the number of single-units with at least a single spike for a given square window (20 or 100ms width), advanced in 1 ms intervals, was calculated. Then the maximal synchronization for non-overlapping windows of the same width was calculated from the summed trace. Thus the largest peak event size possible +/-the window was size was determined for each window. To calculate change in probability spike-times were jittered by a random amount between -1000 and 1000 ms and the probabilities recalculated. For animals with greater than 12 single-units the random assortment was repeated 20 times and the mean probability and jittered probability reported. Binary firing vectors were calculated using the shared Matlab function as described (Okun et al., 2012). For calculation of the Raster-Marginal spikes were swapped between good neurons as well as MUA but only for spikes in the same layer group (L2-4 or L5-6). Calculation of spike-rate correlation followed the method of (Renart et al., 2010). Each single-unit raster was convolved by a normalized kernel that was the sum of a rapid time-window (J, 20ms SD Guassian) and a negative longer window (T, 4xJ). This approximates the effect of jittering the spikes over the same window to remove distorting effects of slow co-modulations in spike-rate. For comparison with truly random firing with locally appropriate spike-rates, jittered spike-trains (+/-1s) were calculated, spike-rate comodulation calculated, and this process was repeated 100 times to generate mean and 95% confidence interval. Population coupling was computed by the method of (Okun et al., 2015). A summed multi-unit raster was created from the MUA and single-units for L2-4 and L5-6 separately, convolved with a Gaussian kernel (10ms SD) which used for spike-triggered average for each single-unit in the corresponding layer group. A normalized spike-triggered rate was made after calculating the Raster Marginal as described above, the peak amplitude of which is used to normalize each animal’s population coupling.

### Statistics

All statistical tests are described in the results along with p-values. P-values below 0.001 are rounded to the nearest decade.

